# Maintenance of variation in virulence and reproduction in populations of an agricultural plant pathogen

**DOI:** 10.1101/2020.04.15.043208

**Authors:** Anik Dutta, Daniel Croll, Bruce A. McDonald, Luke G. Barrett

## Abstract

Genetic diversity within pathogen populations is critically important for predicting pathogen evolution, disease outcomes and prevalence. However, we lack a good understanding of the processes maintaining genetic variation and constraints on pathogen life-history evolution. Here, we analyzed interactions between 12 wheat host genotypes and 145 strains of *Zymoseptoria tritici* from five global populations to investigate the evolution and maintenance of variation in pathogen virulence and reproduction. We found a strong positive correlation between virulence and reproduction, with substantial variation in both traits maintained within each pathogen population. On average, highly virulent isolates exhibited higher fecundity, which might increase transmission potential in agricultural fields planted to homogeneous hosts at a high density. We further showed that pathogen strains with a narrow host range (i.e. specialists) for fecundity were on average less virulent, and those with a broader host range (i.e. generalists) for virulence were on average less fecund on a given specific host. These trade-offs costs associated with host specialization might constrain the directional evolution of virulence and fecundity. We conclude that selection favoring pathogen strains that are virulent across diverse hosts, coupled with selection that maximizes fecundity on specific hosts, may explain the maintenance of these pathogenicity traits within and among pathogen populations.

## Introduction

Plant pathogens typically maintain high intraspecies diversity for key pathogenic traits. These include virulence (defined here as damage caused to the host), host range, and reproduction (Lannou, 2012). Genetic variation underlying phenotypic trait variation (and corresponding resistance traits in their hosts) can be an important determinant of disease epidemiology and can have important consequences for pathogen fitness, host mortality and host reproduction. For example, genetic variation within a pathogen species can facilitate rapid adaptation to control strategies such as fungicide applications and deployment of host resistance (McDonald and Linde, 2002; Zhan *et al*., 2005). However, despite the importance of pathogenicity traits for determining disease incidence, prevalence, and severity, we still lack a clear understanding of how they evolve within populations.

Virulence in plant pathogens can be defined as the degree of damage (e.g., necrosis) and the corresponding fitness reduction in the host following a pathogen infection (Sacristan and Garcia-Arenal, 2008; Barrett *et al*., 2009). Virulence is (at least in part) a direct consequence of host exploitation and is therefore expected to have strong links with pathogen growth and the development of transmission stages needed to infect new hosts (i.e. fecundity). Virulence is thus a key component of pathogen life-history, influencing both the incidence and impact of disease. Populations of plant pathogens typically harbor high levels of genetic variation for both virulence and fecundity (Sacristan and Garcia-Arenal, 2008; Barrett *et al*., 2009). Furthermore, several studies have demonstrated variable expression of these traits according to host and pathogen genetic background, their interaction, and environmental effects in different host-pathogen systems (Pagan *et al*., 2007; Salvaudon *et al*., 2007; Lannou, 2012; Tack *et al*., 2012). However, despite the central importance of these traits to pathogenicity, the processes underlying their evolution and the maintenance of variation within populations are not well understood.

While fecundity is an important life-history trait, pathogen fitness also critically depends upon the transmission of propagules to a new host (Antonovics, 2017). While increased pathogen fecundity can increase potential for transmission, links between virulence, fecundity and transmission are complicated by the fact that transmission also depends on the host. This complexity is captured by the so called ‘trade-off’ theory, which assumes that virulence is an unavoidable consequence of pathogen reproduction within the host (Lenski and May, 1994; Frank, 1996), and predicts that intermediate levels of virulence will maximize transmission because higher virulence may significantly reduce the life expectancy of the infected host (Anderson and May, 1982; Frank, 1996; Leggett *et al*., 2013). However, the trade-off theory has largely been developed within the context of unmanaged host populations (e.g. humans, wild animals etc.). Here, we examine the evolution of virulence and fecundity within populations of an agricultural plant pathogen. There are some key properties specific to agroecosystems that might influence expectations for the evolution of virulence. Unlike natural systems, hosts are homogenous and planted at a high density (allowing frequent physical contact among plants), properties that potentially select for increasingly high virulence and reproduction (McDonald and Stukenbrock, 2016). In addition, farmers replace these high density, homogenous resources every year, meaning that there are potentially few negative consequences associated with increasing levels of host damage. The evolution of virulence in agroecosystems can be further influenced by the pleiotropic effect of genes affecting fungicide resistance in pathogen populations. Extensive fungicide application can impose strong directional selection for resistance, which can be positively correlated with virulence (Yang *et al*., 2013). Thus, it might be predicted that in agricultural settings, directional selection may result in uniformly high levels of virulence and fecundity (Walsh and Blows, 2009; Roff, 2012).

Countering this prediction are frequent reports of high levels of pathogenic diversity within populations of agricultural plant pathogens (Burdon *et al*., 2016). Trade-offs between different components of fitness are often invoked to explain the maintenance of trait variation within species (Stearns, 1989; Thrall *et al*., 2005; Héraudet *et al*., 2008). For host-pathogen interactions, one common prediction is that host specialization will result in the evolution of high levels of fecundity on a restricted subset of hosts (Barrett and Heil, 2012). This outcome implies the existence of trade-offs between the capacity to attack multiple hosts and another component of fitness. A broader host range (i.e. generalism) is expected to increase the number of individual hosts available to infect and lower the risk of extinction should any one host become unavailable, whereas specialization on any individual host (i.e. specialism) comes at the expense of reduced performance on other possible hosts, creating an evolutionary constraint (Kassen, 2002). This specialization trade-off is frequently used to explain why pathogenic traits do not become fixed (i.e. the trade-off maintains trait diversity) in pathogen populations (Brown, 2003). Yet, empirical evidence for costs arising from host specialization are limited (Barrett and Heil, 2012).

*Z. tritici* is the causal agent of septoria tritici blotch, a major fungal disease of wheat (Dean *et al*., 2012; Fones and Gurr, 2015). The pathogen causes necrotic lesions upon infection and produces asexual fruiting bodies called pycnidia within the lesions. It undergoes several cycles of a/sexual reproduction in a growing season (Karisto *et al*., 2018). Lesion development and pycnidia formation were shown to be two different traits, which can vary according to the particular genotypes involved in each host-pathogen interaction (Karisto *et al*., 2018). In this study, we used lesion development and pycnidia formation within lesions as proxies for pathogen virulence and reproduction, respectively. We used a set of 12 wheat host cultivars and 145 *Zymoseptoria tritici* strains to address the following questions: To what extent do key pathogenic life-history traits vary within and among populations of an agricultural plant pathogen? How do pathogen virulence and reproductive traits correlate? What is the impact of spatial structure (i.e. hosts and pathogen populations) on variation in virulence and reproduction? Is there any evidence for a trade-off between specialist and generalist strategies, and if so, what other traits are involved?

## Materials and Methods

### Fungal material

Five genetically different pathogen populations comprising 145 fully sequenced isolates of *Z. tritici* were used in this study. These field populations originated from single wheat fields located in four countries around the world (Zhan *et al*., 2005). The field populations Australia (*n=27*), Switzerland (*n=32*), Israel (*n=30*), and USA (Oregon.R, *n=26*; Oregon.S, *n=30*) were collected in 2001, 1999, 1991 and 1990, respectively. The two populations from Oregon were collected on the same day from two different wheat cultivars, Madsen (Oregon.R) and Stephens (Oregon.S), growing in the same field. All the other populations were sampled from single cultivars. The cultivars Madsen and Stephens were partially resistant and highly susceptible to STB, respectively (Cowger *et al*., 2000). The absence of clones among these isolates and the general absence of pathogen clones within and among populations beyond spatial scales of 1 m was confirmed by previous studies (Linde *et al*., 2002; Zhan *et al*., 2005). After collection, several copies of each isolate were stored and maintained in anhydrous silica gel or 50% glycerol at −80°C.

### Plant material

A set of 12 different wheat hosts were used in this study. This host panel included five landraces (Chinese Spring, 1011, 1204, 4391 and 5254), six commercial varieties (Drifter, Gene, Greina, Runal, Titlis, Toronit) and a back-cross line (ArinaLr34). The 1011, 1204, 4391, and 5254 landraces were selected from a collection of 199 Swiss wheat landraces from the Swiss national gene bank (www.bdn.ch). This panel was screened for resistance to STB (unpublished results) using four fully sequenced *Z. tritici* isolates, namely 3D1, 3D7, 1E4 and 1A5 (Lendenmann *et al*., 2014; Croll *et al*., 2013). The landraces 1011 and 4391 were highly resistant and susceptible, respectively, to all four isolates. The landraces 1204 and 5254 were moderately susceptible to the four isolates. The remaining hosts have diverse genetic backgrounds and were previously used to identify infectivity (i.e. a/virulence) factors in *Z. tritici* (Hartmann *et al*., 2017; Zhong *et al*., 2017; Meile *et al*., 2018; Stewart *et al*., 2018). The seeds of Gene and ArinaLr34 were provided by Christopher Mundt (Oregon State University) and Simon Krattinger (KAUST), respectively. The seeds of other hosts were obtained from DSP Ltd. (Delley, Switzerland).

### Preparation of fungal inoculum

The isolates were regenerated from long term glycerol storage by adding 40μl of concentrated spore suspension into 100 ml Erlenmeyer flasks containing 50ml of yeast sucrose broth (YSB, 10 g/L sucrose, 10 g/L yeast extract amended with 50 μg/ml kanamycin sulfate to control other microbial growth). The flasks were kept at 18°C on a continuous shaker at 120 rpm to produce blastospores. Blastospores were harvested after 4-7 days of growth by filtering the liquid media through two layers of sterile cheesecloth. Blastospore pellets were collected by centrifugation (1575 g, 15 minutes, 4°C), washed with sterile water to eliminate any residual growth media and re-suspended in sterile water for subsequent procedures. The spores were counted and adjusted to a final concentration of 5×10^6^ spores/ml using KOVA counting slides (Hycor Biomedical, Inc., Garden Grove, CA, USA). The spore suspension of each isolate was stored at −20°C until inoculation for between 1-21 days.

### Phenotyping and data collection

Due to greenhouse space limitations, the whole experiment was divided into two phases consisting of a combination of 6 hosts × 145 isolates in each phase. Three seeds of each host were sown in an individual square pot filled with peat substrate Jiffy GO PP7 (Jiffy Products International, Moerdijk, the Netherlands). Six pots were placed on a tray in a 2×3 array. All the trays were kept in a greenhouse at 22°C (day) and 18°C (night) with 70% relative humidity (RH) and 16-h photoperiod. Plants were inoculated after developing a fully expanded second leaf, at 14 days after sowing. The third leaf and subsequent leaves were trimmed before inoculation and new leaves were trimmed until data collection to facilitate a more uniform distribution of spores and better light penetration onto the inoculated leaf. Before the inoculation day, the spore suspension of each isolate was thawed on ice and kept at 4°C overnight. The final volume of each spore suspension used for inoculation was adjusted to 20 ml by adding sterile water supplemented with 0.1% of the Tween 20 surfactant. Each tray containing six hosts was inoculated with an airbrush spray gun (1A Profi Handels GmbH, Wiesbaden, Germany) uniformly until run-off. The spraying was done in a confined area to minimize any chance of cross-contamination. The trays were covered with plastic bags after inoculation to provide 100% RH and transferred into a greenhouse chamber. The entire procedure was repeated three times over three consecutive weeks to generate three independent biological replicates for both phases of the experiment. Spraying of all 145 isolates was performed on a single day. For the landraces 1011, 5254 and 1204, 4391, only one and two plants, respectively, were inoculated in each replicate due to limited seed availability.

Plastic bags were removed three days post-inoculation (dpi). Environmental conditions otherwise remained identical. To facilitate comparisons among hosts, leaves were harvested between 19-26 dpi because different hosts developed symptoms at different rates. All leaves from each host×isolate combination were collected on the same day. Each second leaf was excised and mounted on A4 paper for scanning as described previously (Karisto *et al*., 2018). Each A4 sheet containing eight inoculated leaves was scanned using a flatbed scanner (CanoScan LiDE 220) for automated image analysis (AIA; Karisto *et al*., 2018). The AIA provided quantitative data on the amount of lesion area caused by the fungus and pycnidia density within the lesions on each leaf. As previously described in Karisto *et al*., (2018), lesion area corresponds to host damage and was used as a proxy for virulence, whereas pycnidia density is a direct measure of pathogen asexual reproduction as pycnidia produce the spores that are eventually transmitted through rain splash or direct contact to neighboring plants.

### Data analysis

Before analysis, all phenotypic values were log-transformed to fulfill the normality assumptions of ANOVA based on the residual distribution. The log-transformed values were used for all analyses. A combined analysis of variance (ANOVA) was performed to estimate the effect of different factors on the two traits. The following linear model was implemented using the *lm()* function from the package lme4 (Bates *et al*., 2014) in R (R core team, 2019): *Virulence/Reproduction∼Host+Population+Isolate:Population+Host:Population+ Host:Isolate:Population+Replication+Error*

All factors were treated as fixed effects and isolates were nested within each population. The least-square mean (LSmean) for each host×isolate combination was extracted by using the function “emmeans” from the package emmeans (Lenth, 2018). Using the LSmean from each host, we computed the global mean for each isolate for virulence and reproduction. Posthoc multiple comparisons of LSmeans of the host and pathogen population interactions were performed by using Tukey’s HSD test at the α=0.05 significance level.

We used the coefficient of variation (CV = standard deviation/mean) of LSmeans for each isolate among 12 hosts as a metric for host specificity (Poissot *et al*., 2012; Caseys *et al*., 2019). Several methods including CV have been proposed for measuring specificity at both individual and species levels (Bolnic *et al*., 2002; Poissot *et al*., 2012). However, each of these has limitations depending on the type of data. While the majority of the specificity metrics assume discrete data, very few are available for continuous data. In our dataset, the use of host resources was more evenly distributed. The majority of isolates readily infected the majority of susceptible hosts, while the degree of virulence and the amount of reproduction varied among the majority of them. Hence we favored the use of CV, which facilitates estimation of the degree of specialism and generalism among the isolates using precise quantitative phenotypic data.

To test for trade-offs, we performed Pearson’s correlation analysis between virulence and reproduction, using the overall mean across 12 hosts. Furthermore, we tested the shape of the correlation curve by fitting a polynomial model *(Reproduction∼poly (Virulence, 2))*. We were interested in determining how the specificity of each isolate affects its mean virulence and reproduction. Therefore, we performed correlations between mean virulence and the specificity of reproduction as well as between mean reproduction and the specificity of virulence. All analyses were performed in R-studio.

## Results

### Determinants of quantitative variation in pathogen virulence and reproduction

We obtained quantitative data from 11’019 inoculated leaves generated from the interactions between 12 hosts and 145 pathogen strains using AIA to investigate how variation in pathogen life-history traits are maintained. Lesion area and pycnidia density within the lesion area were used as proxies of virulence and reproduction, respectively. Isolates displayed a wide range of phenotypic responses, exhibiting a quantitative distribution in virulence and reproduction on each host (Supplemental figs. 1A and 1B). Using the non-transformed data, virulence ranged from 0 to 100% with an overall mean of 61.4 ± 0.4%. The host 1011 showed the highest resistance on average, followed by Gene and Toronit (Supplemental Table 1). Mean reproduction across 12 hosts ranged from 0 to 671 pycnidia per cm^2^ of lesion with an overall mean of 17.8 ± 0.4. The isolates were on average highly virulent and fecund on the most susceptible host 4391.

We applied a linear model to test which factors contributed to the observed quantitative variation in virulence and reproduction. There was significant variation (*P* < 2e^−16^) among the hosts, populations, and isolates for both the traits (Table 1). Host identity strongly influenced the observed variation in reproduction (F = 835.53, *P* < 2e^−16^) and virulence (F = 232.47, *P* < 2e^−16^). We obtained a highly significant interaction effect between hosts and isolates nested within each population for virulence (F = 2.11, *P* < 2e^−16^) and reproduction (F = 2.68, *P* < 2e^− 16^). This significant interaction effect indicates the occurrence of host specificity among the isolates for both traits. Multiple comparisons among the populations on each host revealed high variability and changes in ranking among the populations for both traits (Figs. 1A and 1B). The two populations from Oregon had the highest overall mean virulence. The Israeli population exhibited the highest reproduction, followed by Oregon.R and Switzerland.

**Table 1.**
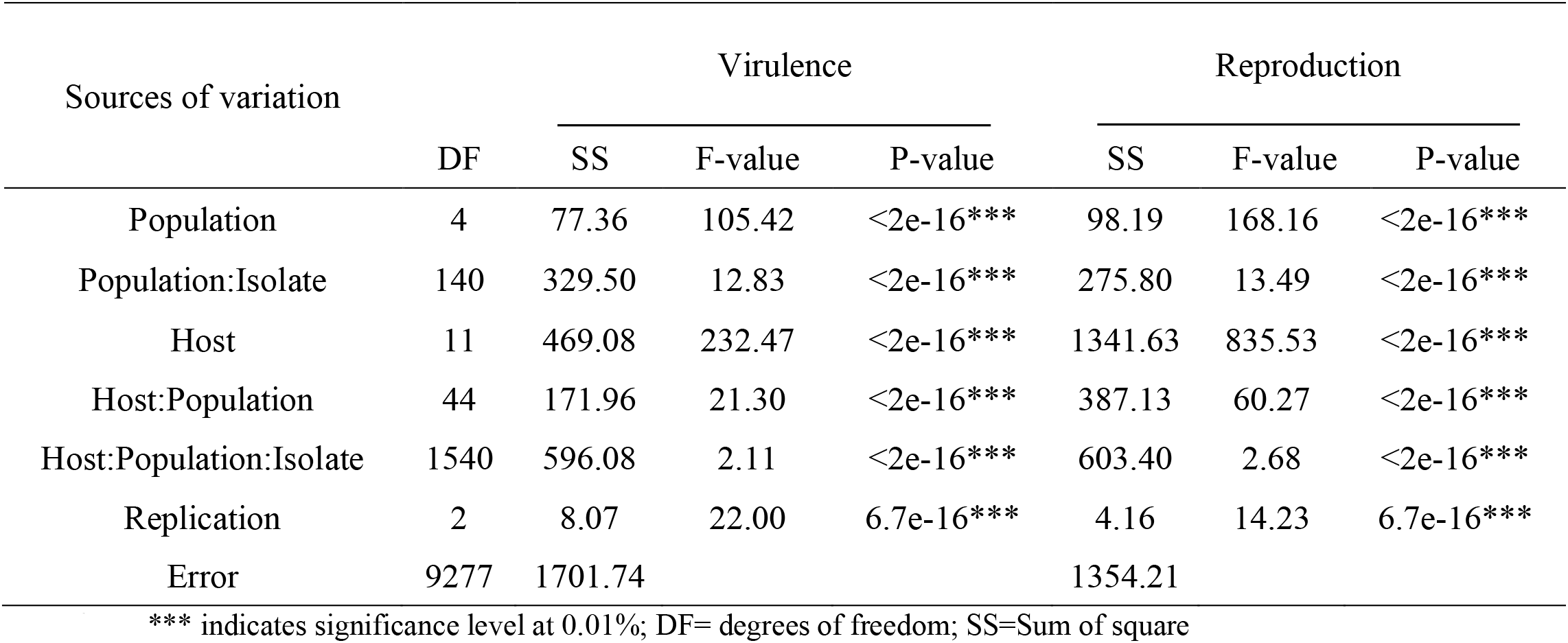
Analysis of Variance (ANOVA) showing the effects of hosts, populations, isolates and the respective interactions on virulence (amount of necrotic lesion area) and reproduction (pycnidia density within lesions) among 145 *Zymoseptoria tritici* isolates from five populations across 12 hosts.

**Figure 1.**
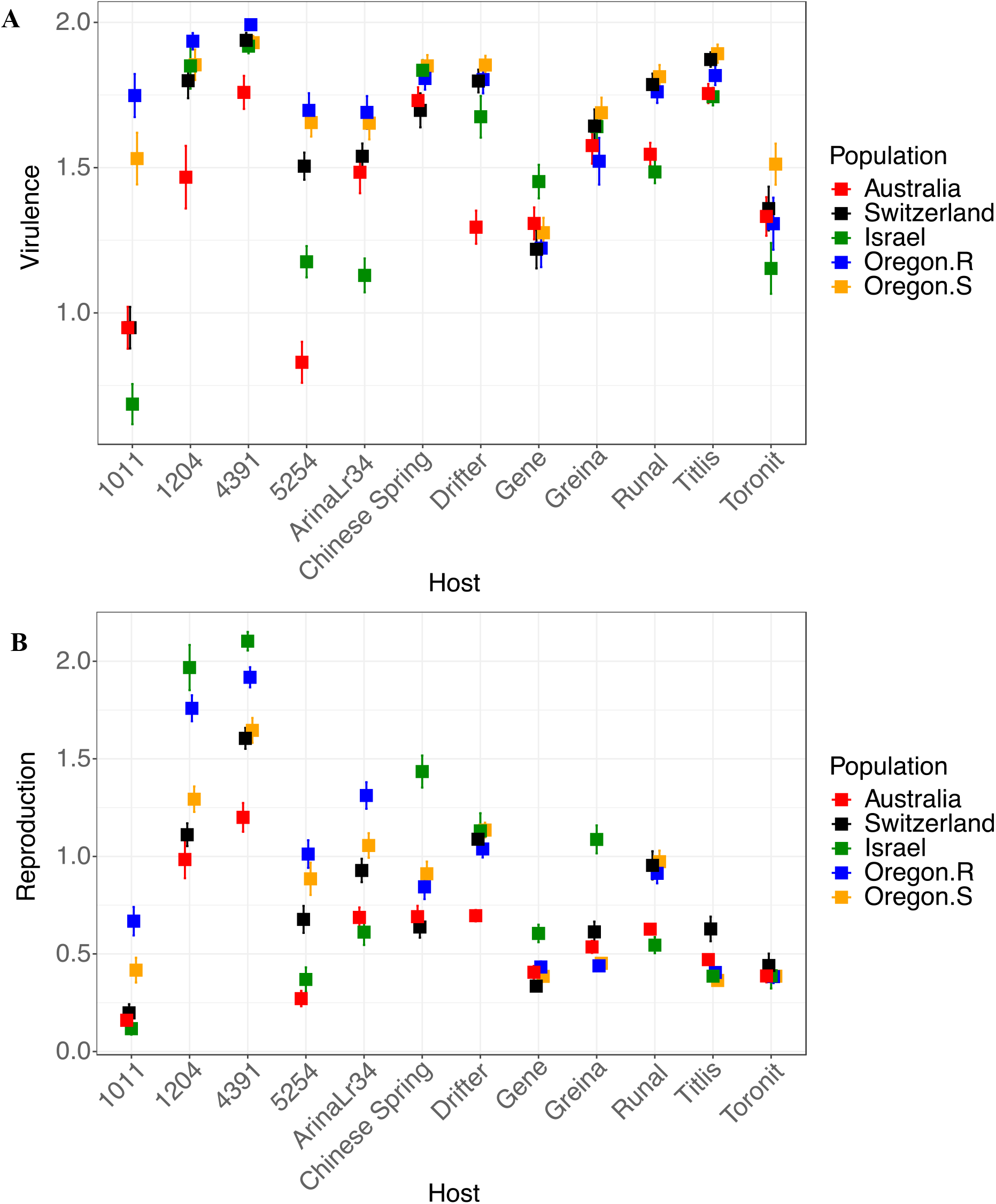
Multiple comparisons for (A) virulence (amount of necrotic lesion area) and (B) reproduction (pycnidia density within lesions) among the five *Zymoseptoria tritici* populations on 12 wheat hosts. Data were log-transformed.

### Absence of a trade-off between pathogen virulence and reproduction

We performed Pearson’s correlation between virulence and reproduction to determine whether or not there is a trade-off between these traits. The overall mean values of each isolate across 12 hosts were used in this analysis. Overall, we detected a significant positive correlation (*r* = 0.62, *P* < 2.2e^−16^; Fig. 2A) between the two traits. This indicates that highly virulent isolates also had high levels of fecundity. However, the polynomial regression did not show evidence for any saturating point (results not shown) on the curve, indicating an increasing trend for both traits. The positive correlation was consistent within each population, although the strength of the correlation within each population varied considerably, with the ISR population showing the highest correlation coefficient (*r* = 0.49 to 0.85, *P* = 0.0045 to 3.5e^−09^; Fig. 2B). The variation in the correlation indicated that individual isolates might differ in their strategy to exploit certain hosts.

**Figure 2.**
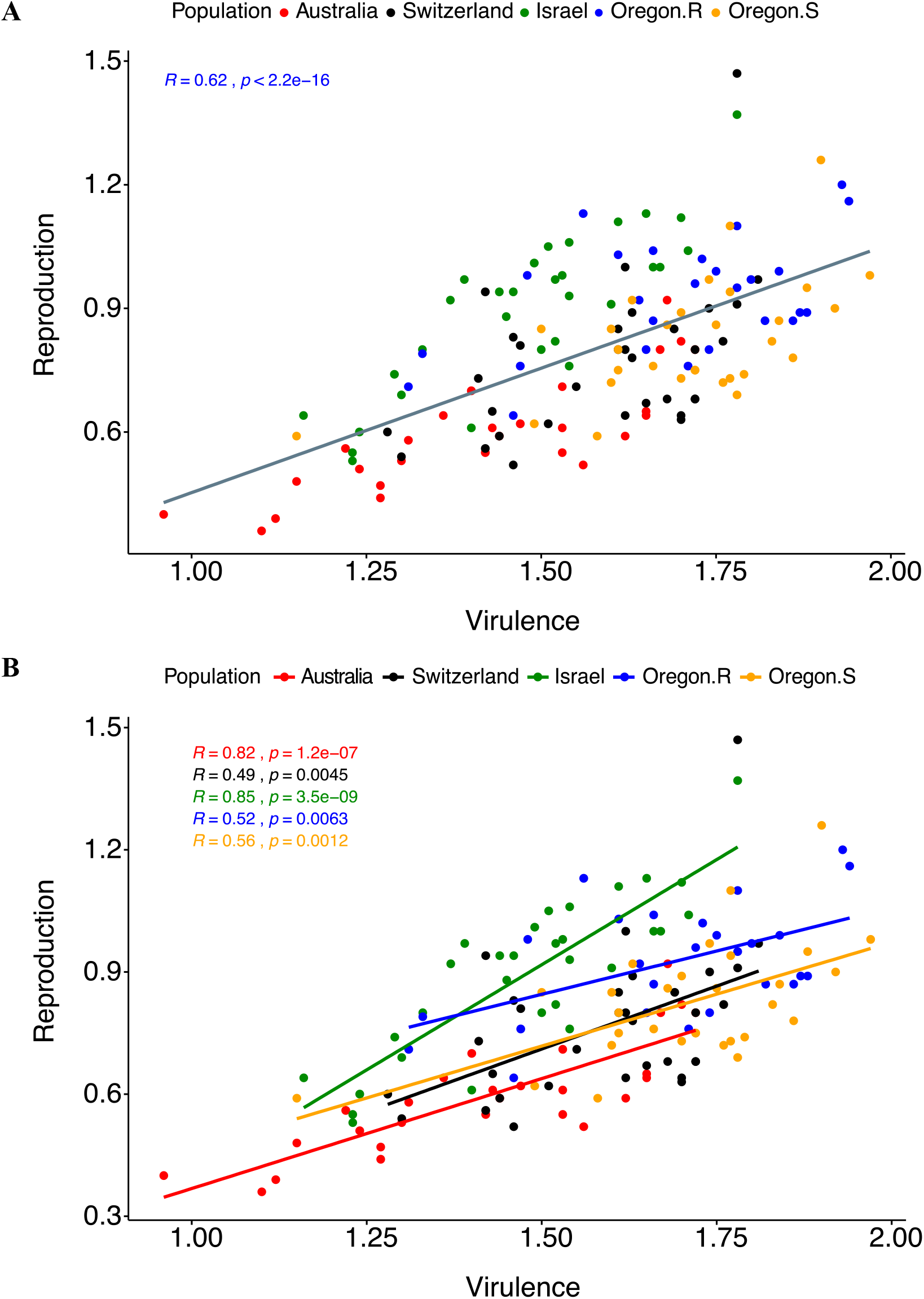
Correlation between virulence (amount of necrotic lesion area) and reproduction (pycnidia density within lesions; A) overall and (B) within each population, among 145 *Zymoseptoria tritici* isolates from five populations. Each point represents the overall.

### Host specialization reduces the mean trait performance

We investigated the influence of host specialization on the overall virulence and reproduction and the maintenance of genetic diversity in the field. Using CV as a specificity metric, we detected a significant negative correlation (*r* = −0.60, *P* = 8.5e^−16^, Fig. 3A) between specialization for reproduction and mean virulence. This pattern indicates that isolates that are generalists for reproduction have higher mean virulence across the 12 hosts. We also found a significant negative correlation (*r* = −0.42, *P* = 1.3e^−07^, Fig. 3B) between specialization for virulence and mean reproduction, indicating that isolates that exhibit host specialization for virulence have an overall lower rate of reproduction across the 12 hosts. It is evident that most of the isolates from the populations in Oregon and Israel are generalists or specialists, respectively, for both traits (Figs. 3A & 3B). Importantly, the positive correlation (*r* = 0.46, *P* = 6.8e^−09^, Fig. 4) between specialization for reproduction and maximum reproduction indicate that there is a benefit associated with being a specialist.

**Figure 3.**
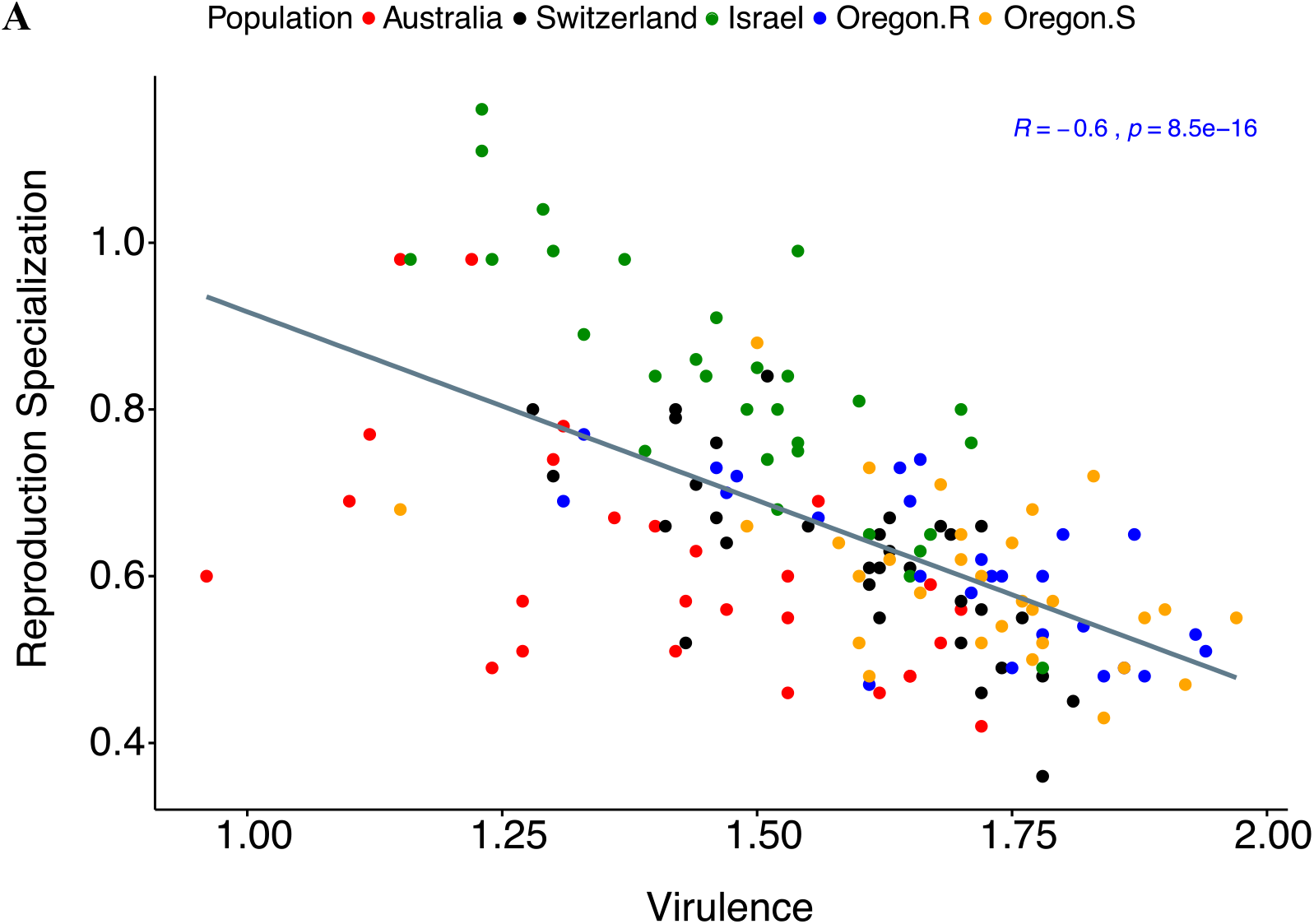
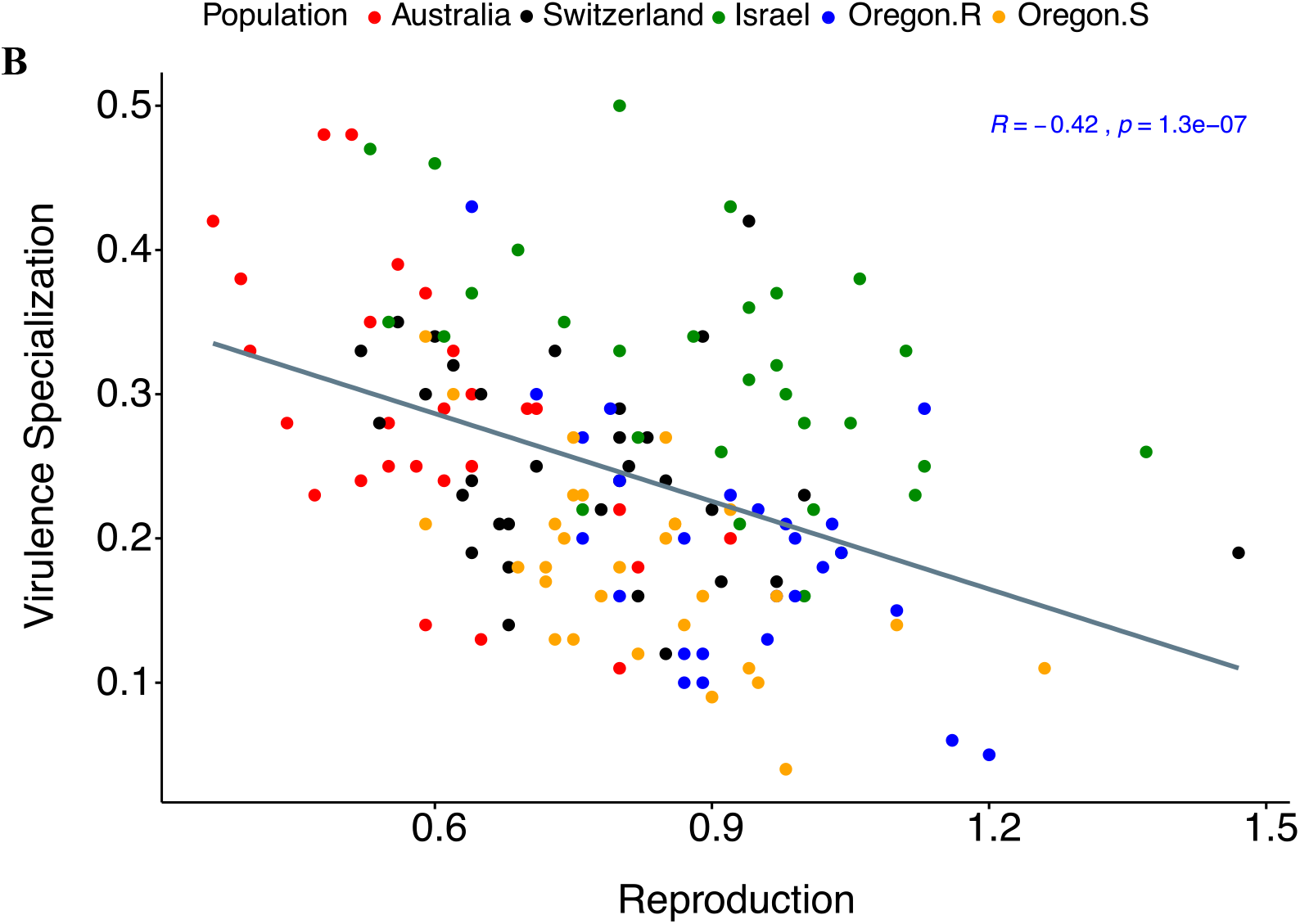
Correlation between (A) overall mean virulence (amount of necrotic lesion area) and reproduction specialization, (B) overall reproduction (pycnidia density within lesions) and virulence specialization among 145 *Zymoseptoria tritici* isolates from five populations. Specialization represents the estimates of coefficient of variation of means across 12 hosts for each trait. Higher specialization indicates preference for specific hosts to maximize trait performance. Data were log-transformed.

**Figure 4.**
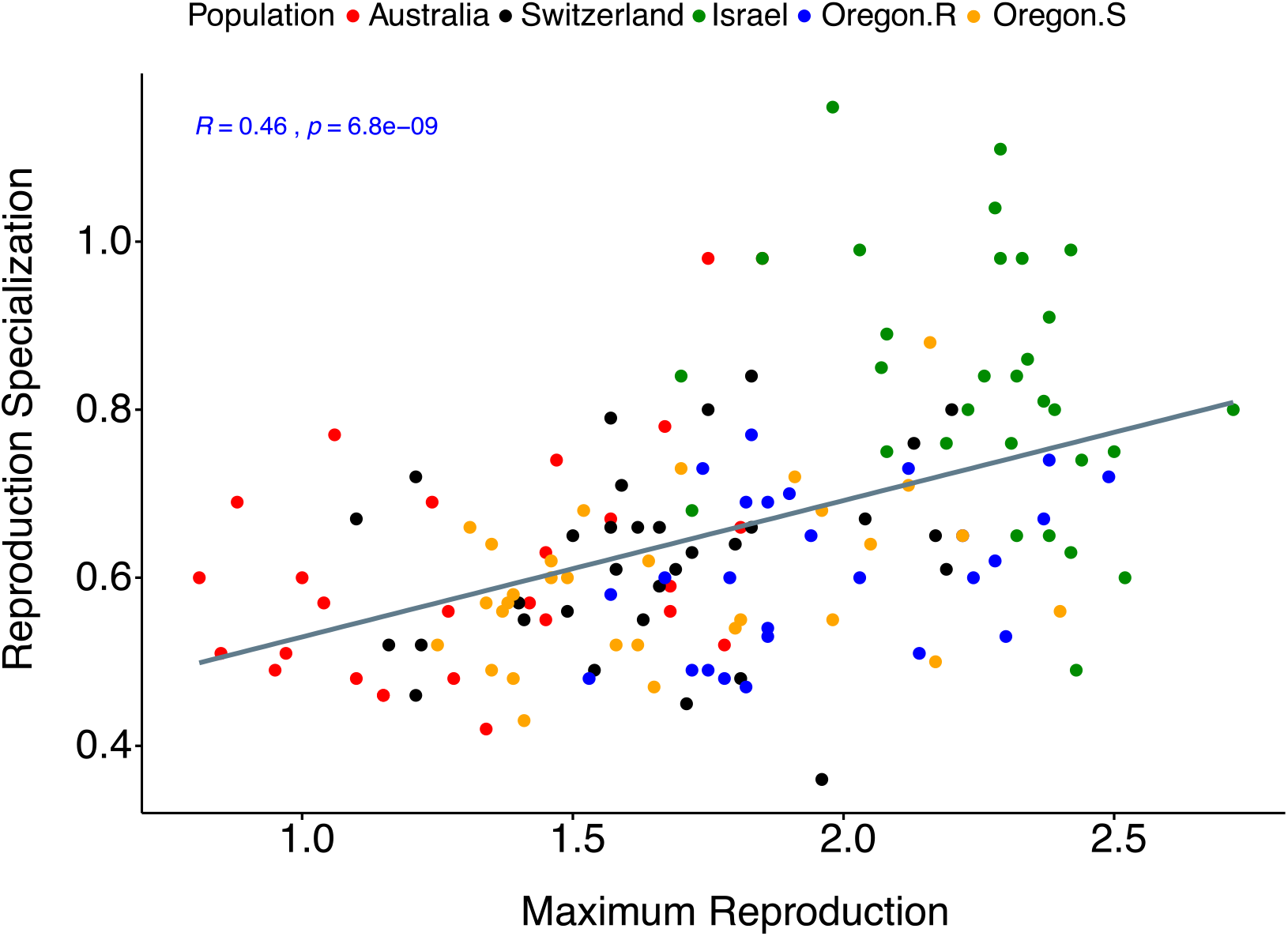
Correlation between maximum reproduction (maximum pycnidia density within the lesion area produced by each isolate across 12 hosts) and reproduction specialization (measured as the coefficient of variation of means for pycnidia density within lesions across 12 hosts) among 145 *Zymoseptoria tritici* isolates from five populations. Data were log-transformed.

## Discussion

Understanding the processes that maintain the variation in pathogenicity is of central importance to predict pathogen evolution and understand disease dynamics (Burdon and Thrall, 2008). Using a panel of 12 diverse hosts and 145 pathogen strains, we demonstrated that individual estimates of virulence and reproduction varied quantitatively, and that host genotype and interactions with pathogen genotype are major contributors to the observed variation in virulence and reproduction. We observed a strong positive correlation between the two traits with substantial variation within each population. Furthermore, the data show a continuum of host specialization among *Z. tritici* isolates, where isolates ranged between high and low fecundity depending on the specific host infected. This may constrain directional evolution of both virulence and fecundity and contribute to the maintenance of trait variation.

### Host diversity as a key determinant of pathogenicity trait variation

Infection outcomes in many pathosystems are determined by genetic interactions among hosts and pathogens (e.g. Pagan *et al*., 2007; Pariaud *et al*., 2009; Lannou, 2012; Tack *et al*., 2012). Earlier models of pathogen evolution often assumed that variation in infectivity is solely controlled by pathogen strains (Restif and Koella, 2003). Here, in addition to the significant host-isolate interactions, the effect of host cultivar on reproduction was almost 3-fold higher than that observed for virulence, indicating that host genetic background strongly influences resource allocation for reproduction following pathogen colonization (Vale *et al*., 2011; Attisano *et al*., 2012). This result is consistent with previous studies, which show that even after successful infection, host immune response, nutrient availability, and quantitative resistance may hamper pathogen reproduction (Karisto *et al*., 2018). The high level of dependency on host genotype for reproductive fitness is consistent with results demonstrating that different isolates maximize reproductive fitness on specific hosts and that host specific patterns of fecundity are potentially important for the maintenance of pathogen diversity (see discussion below).

### Evolution of pathogen virulence and fecundity in agroecosystems

Here we show that highly virulent isolates of *Z. tritici* on average exhibit greater reproductive potential. This likely reflects general links between pathogen induced necrosis and nutrient release in this system (Zhan *et al*., 2005; Kelm *et al*., 2012). We did not detect any evidence for a saturation point on the correlation curve generated from our dataset, suggesting that both traits follow a continuous range. On a susceptible host, high levels of fecundity and virulence are likely to be advantageous because highly virulent isolates can outcompete co-infecting strains (Zhan *et al*., 2015; McDonald and Stukenbrock, 2016). However, despite this seeming advantage, highly virulent and fecund isolates did not dominate any population. Rather, we observed that considerable variation for both virulence and fecundity is maintained within field populations.

Trade-offs between transmission and virulence have been predicted to contribute to the maintenance of variation for virulence. This reflects the assumption that increased virulence decreases the longevity of the infected host so that the transmission period decreases, creating a trade-off (Ebert and Bull, 2003; de Roode *et al*., 2008). In this agricultural ecosystem (characterized by high density monocropping with relatively homogeneous hosts), higher reproduction (and higher virulence) may be predicted to increase transmission potential as the reproductive units (pycnidia) contain spores that are dispersed by rain-splash throughout the field, and hosts are replaced by farmers each growing season. However, it should be noted that although fecundity can influence transmission, they are not synonymous. Under natural field conditions, transmission depends on other factors such as the quantity, viability, and infection efficiency of spores, as well as climatic conditions and, most importantly, the availability of a susceptible host. How these traits and outcomes interact to determine virulence evolution is not well understood.

We found considerable variation among populations in overall levels of virulence and reproduction on a diverse set of hosts. We also found a strong correlation between these traits within populations. These results suggest that selection for these traits can differ among populations. One possible explanation for these findings is that the differential patterns of virulence and reproduction reflect specific patterns of adaptation to the most common host genotypes planted in the different wheat fields from which these pathogens were sampled. However, the Swiss population displayed relatively low levels of virulence and fecundity, which was surprising considering that the bulk of the tested hosts originated from Swiss breeding programs. It is possible that the intensive breeding for STB resistance and frequent introductions of new varieties from neighboring countries (Fossati and Brabant, 2003; Brabant *et al*., 2006) in Switzerland may have hampered selection for higher trait performance and adaptation to a specific host. Thus, imposing diversifying selection on local pathogen populations by recurrent changing of wheat cultivars and specific resistance traits might be a useful strategy to disrupt evolution towards increased virulence and fecundity (Burdon *et al*., 2014). Furthermore, in addition to quantitative resistance to *Z. tritici* (Yates *et al*., 2019), wheat varieties across the world often share common major resistance genes (Brown *et al*., 2015), which can be overcome by convergent evolution generating similar virulent mutations independently in different geographical populations, even in the absence of gene flow among populations (Croll and McDonald, 2017). These processes could explain the observed higher performance of the Israeli and Oregon populations on Swiss hosts.

### Role of host specialization in the evolution of pathogenicity trait variation

Does host specialization limit the emergence of “super-pathogen” strains combining higher virulence, higher fecundity, and a broader host range? Here, all the hosts exhibited a wide spectrum of resistance and susceptibility towards pathogen virulence and reproduction. For example, the wheat genotypes 1011, Gene and Toronit showed greater resistance to virulence and suppressed reproduction of some isolates, while other wheat genotypes were moderately to highly susceptible to virulence and enabled higher reproduction of some isolates. Therefore, we assessed patterns of host specificity by examining patterns of quantitative variation for virulence and reproduction among the many host-isolate interactions in our experiment. We found significant negative correlations between overall virulence and reproduction specificity, indicating that isolates with high host specificity for reproduction (i.e. higher pycnidia production on some hosts and a smaller number of pycnidia on other hosts) were on average less virulent across all hosts (i.e. produced a lower average lesion area). This observation is consistent with the hypothesis that increased specificity for fecundity results in decreased virulence (Kirchner and Roy, 2002). This pattern further indicates a cost associated with host specialization, which is suggested to result from antagonistic pleiotropy or non-overlapping loci controlling trait performance on different hosts (Kawecki, 1994; Legros and Koella, 2010; Hartmann *et al*., 2017).

The observed positive relationship between specialization for reproduction and maximum reproduction suggests that isolates following a generalist strategy fail to maximize their reproductive fitness on a specific host, thus invoking the principle of “jack of all trades-master of none” (Remold, 2012). This trade-off could explain why specialist isolates are maintained within populations and why ‘super-pathogen’ strains do not dominate (Kassen, 2002). This is consistent with the results reported by Thrall and Burdon (2003), where generalist isolates with a broader host range had lower overall fecundity. Such specificity for reproduction may be advantageous under scenarios involving host heterogeneity, competition during multiple infection, and disruptive selection (Jaenike, 1990; Barrett *et al*., 2009).

The precise scenario(s) generating heterogeneity, disruptive selection and promoting the maintenance of variation for host specialization within and among populations remain unknown. Specialists could outperform generalists on a given host (Garamszegi, 2006; Romero and Elena, 2008) because generalists suffer from unequal selection pressure imposed by different hosts (González *et al*., 2019). For example, in a wheat field planted to a susceptible cultivar (typically grown in a monoculture), being highly specialized in reproduction could be beneficial because competition among isolates can favour selection for higher reproduction instead of broader host range. In contrast, strains with high average levels of fecundity across different host genotypes may be expected to have an advantage in environments with higher levels of host genetic heterogeneity (May and Anderson, 1983; Frank, 1996). For example, in areas where multiple cultivars are planted in relatively close proximity, generalist pathotypes may have an advantage. Indeed, the maintenance of variation may reflect conflicts between selection for fecundity and transmission within fields planted to a single susceptible cultivar and selection for ability to infect multiple cultivars planted across an agricultural landscape. In our case, whether the diversity of wheat host genotypes present within agricultural landscapes is alone sufficient to explain the maintenance of generalist pathotypes given the presumed advantage to specialists remains an interesting but open question.

## Conclusion

Here, we report how different processes regulate life-history trait variation, which ultimately improves our understanding of pathogen evolution and disease dynamics. Host diversity and differential quantitative interactions with pathogen strains are likely key determinants of variation in virulence and reproduction. Trade-offs for reproduction encountered by specialist vs generalist isolates reinforce the general importance of costs in maintaining pathogenicity trait variation. These mechanisms prevent the fixation of super-pathogen strains in pathogen populations while indicating that diversifying the host in agricultural fields might be a useful strategy to decelerate virulence evolution. Many approaches can be used to introduce dynamic diversity into agricultural ecosystems and reduce selection for increased virulence (McDonald, 2014). A low tech approach that was shown in independent, replicated, field experiments to impose diversifying selection on populations of three cereal pathogens is growing cultivar mixtures (Zhan and McDonald, 2013). Implementing such dynamic diversity at the field scale may impose such trade-offs on generalist isolates that lowers the ability to maximize reproduction on any particular host (McDonald, 2014). We conclude that context-dependent selective forces operating on pathogen populations located in different geographic locations will likely play an ongoing role in contributing to the maintenance of variation in virulence and reproduction in this and other pathosystems. While increased reproduction is likely to provide more opportunities for transmission and virulence evolution in the environments that typify much of modern agriculture, we still lack a deeper understanding of the impact of other life-history traits that have an impact on epidemiology and disease dynamics, such as latent period, spore quantity and infection efficiency.

## Data availability

Raw data used for this study are available at: https://doi.org/10.5061/dryad.j3tx95x9m.

## Author contributions

LGB conceived the idea of the research. AD conducted the experiment, collected, and analyzed the phenotypic data, wrote the manuscript with LGB. BAM and DC provided funding and corrected the manuscript. All authors approved the final version of the manuscript.

## Acknowledgments

This work was supported by the Swiss Federal Office for Agriculture (BLW) in the framework of NAP-PGREL (national plan of action for the conservation and sustainable utilization of plant genetic resources) Project Nr. 627000640. Peter Thrall read an earlier version and made useful comments and suggestions that improved the manuscript.

