## Supplemental Table 1, Supplemental Fig. 1 for "Maintenance of variation in virulence and reproduction in populations of an agricultural plant pathogen"

**Supplemental Table 1**. Summary statistics of virulence (amount of necrotic lesion area) and reproduction (pycnidia density within lesions) in each host and population from 145 *Zymoseptoria tritici* isolates using the non-transformed data.

| | | Virulence | | | Reproduction | | |
|---|---|---|---|---|---|---|---|
| | | Mean | Range | SE* | Mean | Range | SE |
| Host | 1011 | 36.8 | 0-100 | 2.5 | 3.2 | 0-79 | 0.6 |
| | 1204 | 77.3 | 0-100 | 1.4 | 70.0 | 0-671 | 4.3 |
| | 4391 | 85.7 | 2.7-100 | 1.1 | 87.8 | 0-588 | 3.9 |
| | 5254 | 49.9 | 0-100 | 2.1 | 14.1 | 0-332 | 1.7 |
| | ArinaLr34 | 57.9 | 0-100 | 1.4 | 20.7 | 0-382 | 1.4 |
| | Chinese Spring | 78.2 | 0-100 | 1.0 | 22.5 | 0-538 | 1.3 |
| | Drifter | 68.3 | 0-100 | 1.1 | 20.0 | 0-418 | 1.0 |
| | Gene | 38.9 | 0-100 | 1.0 | 4.1 | 0-200 | 0.3 |
| | Greina | 60.7 | 0-100 | 1.0 | 8.1 | 0-205 | 0.5 |
| | Runal | 59.7 | 0-100 | 0.9 | 10.9 | 0-220 | 0.5 |
| | Titlis | 75.5 | 0-100 | 0.8 | 3.5 | 0-92.3 | 0.3 |
| | Toronit | 42.2 | 0-100 | 1.1 | 3.1 | 0-74.8 | 0.2 |
| Population | Australia | 50.1 | 0-100 | 0.8 | 6.72 | 0-275 | 0.4 |
| | Israel | 56.2 | 0-100 | 0.8 | 33.9 | 0-671 | 1.5 |
| | Oregon.R | 70.0 | 0-100 | 0.7 | 14.2 | 0-426 | 0.6 |
| | Oregon.S | 66.5 | 0-100 | 0.8 | 20.7 | 0-588 | 1.1 |
| | Switzerland | 63.8 | 0-100 | 0.8 | 13.1 | 0-382 | 0.6 |

*SE= Standard Error

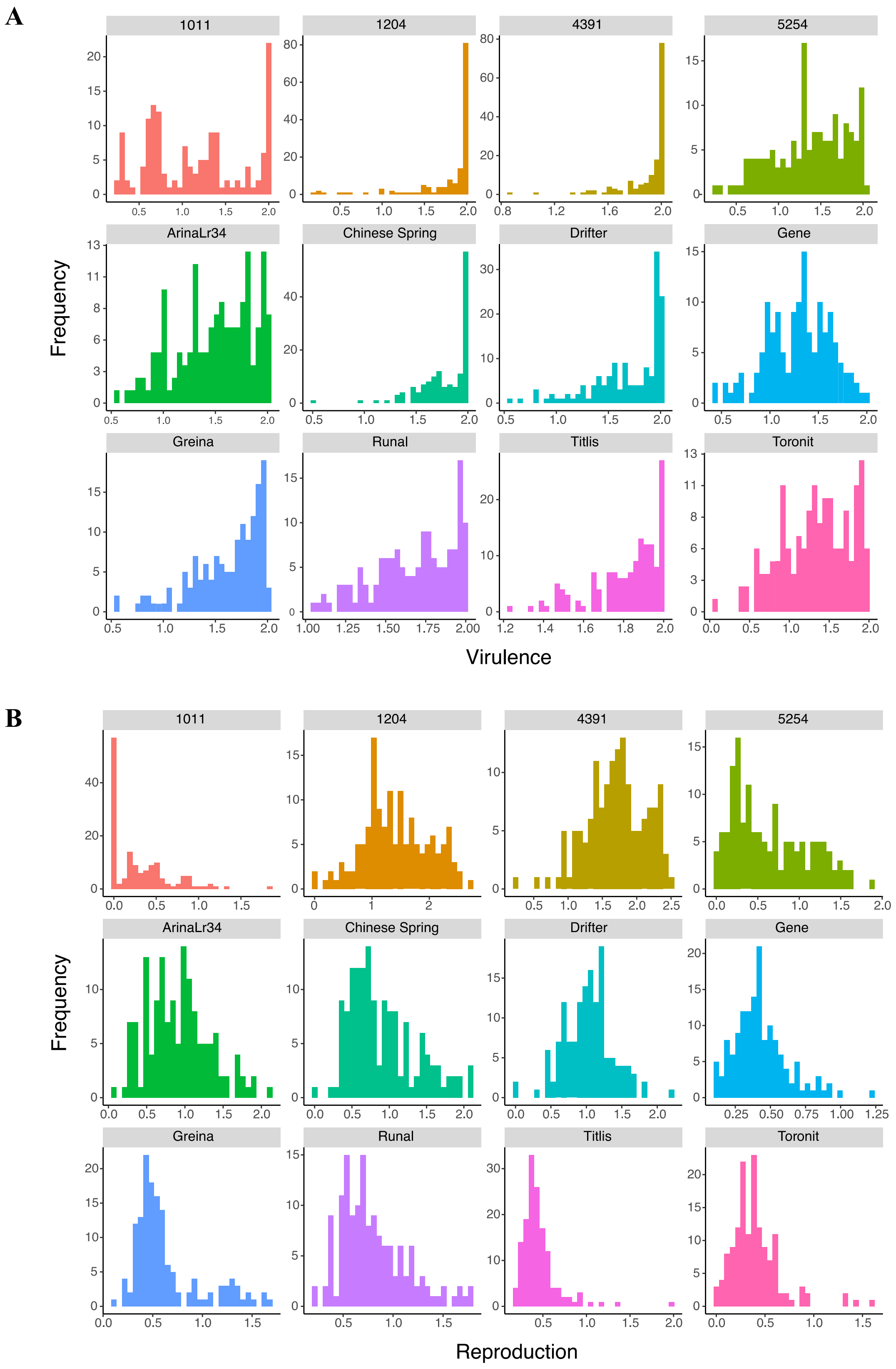


**Supplemental Figure 1**. Frequency distribution of (A) virulence (amount of necrotic lesion area) and (B) reproduction (pycnidia density within lesions) among 145 *Zymoseptoria tritici* isolates from five populations in 12 hosts. Data were log-transformed.
